# IGF-1 Acts through Kiss1-expressing Cells to Influence Metabolism and Reproduction

**DOI:** 10.1101/2024.07.02.601722

**Authors:** Mengjie Wang, Seamus M. Pugh, Judy Daboul, David Miller, Yong Xu, Jennifer W. Hill

## Abstract

**Objective:** Kisspeptin, encoded by the *Kiss1* gene, ties puberty and fertility to energy status; however, the metabolic factors that control *Kiss1*-expressing cells need to be clarified.

**Methods:** To evaluate the impact of IGF-1 on the metabolic and reproductive functions of kisspeptin producing cells, we created mice with IGF-1 receptor deletion driven by the *Kiss1* promoter (IGF1R^Kiss1^ mice). Previous studies have shown IGF-1 and insulin can bind to each other’s receptor, permitting IGF-1 signaling in the absence of IGF1R. Therefore, we also generated mice with simultaneous deletion of the IGF1R and insulin receptor (IR) in *Kiss1*-expressing cells (IGF1R/IR^Kiss1^ mice).

**Results:** Loss of IGF1R in *Kiss1* cells caused stunted body length. In addition, female IGF1R^Kiss1^ mice displayed lower body weight and food intake plus higher energy expenditure and physical activity. This phenotype was linked to higher proopiomelanocortin (POMC) expression and heightened brown adipose tissue (BAT) thermogenesis. Male IGF1R^Kiss1^ mice had mild changes in metabolic functions. Moreover, IGF1R^Kiss1^ mice of both sexes experienced delayed puberty. Notably, male IGF1R^Kiss1^ mice had impaired adulthood fertility accompanied by lower gonadotropin and testosterone levels. Thus, IGF1R in *Kiss1*-expressing cells impacts metabolism and reproduction in a sex-specific manner. IGF1R/IR^Kiss1^ mice had higher fat mass and glucose intolerance, suggesting IGF1R and IR in *Kiss1*-expressing cells together regulate body composition and glucose homeostasis.

**Conclusions:** Overall, our study shows that IGF1R and IR in *Kiss1* have cooperative roles in body length, metabolism, and reproduction.

## 1. INTRODUCTION

Kisspeptin (encoded by the *Kiss1* gene) is a critical regulator of both puberty onset and fertility in mammals with expression throughout the reproductive axis [1–3]. In the brain, kisspeptin exerts its effects on the hypothalamic-pituitary-gonadal (HPG) axis by stimulating gonadotropin-releasing hormone (GnRH) neuron activity [4; 5]. Reproduction requires adequate energy stores [6]. *Kiss1*-expressing neurons have been previously hypothesized to serve as a primary transmitter of metabolic signals from the periphery to GnRH neurons [6–11]. Studies have documented a clear impact of nutrition or metabolic stress on *Kiss1* expression in the hypothalamus [9; 12-14]. For example, fasting mice displayed reduced hypothalamic *Kiss1* mRNA levels, which preceded a decline in GnRH expression [12]. Chronic subnutrition during puberty reduced *Kiss1* mRNA levels in female rats [14]. Furthermore, repeated central injections of kisspeptin restored pubertal progression to female rats subjected to chronic subnutrition [14]. Thus, *Kiss1*-expressing neurons can coordinate reproduction with metabolic status.

*Kiss1*-expressing neurons in mice are mainly located in two regions of the hypothalamus, one in the arcuate nucleus of hypothalamus (ARH) and the other in the rostral periventricular area of the third ventricle (RP3V) [15–17]. Recently, *Kiss1*-expressing neurons of the ARH have been shown to be involved in diet-induced obesity in several animal models [18–20]. Female mice with deletion of the receptor for the gut peptide ghrelin in *Kiss1*-expressing cells were resistant to body weight gain on a high fat diet (HFD) [18]. In addition, lack of the endoplasmic reticulum calcium-sensing protein stromal interaction molecule 1 (STIM1) in *Kiss1* neurons of the ARH protected ovariectomized female mice from developing obesity and glucose intolerance with HFD. The mechanisms underlying how *Kiss1*-expressing cells regulate metabolism are beginning to be uncovered. Optogenetic activation of ARH *Kiss1* neurons evokes glutamatergic excitatory postsynaptic currents in arcuate proopiomelanocortin (POMC) and agouti-related peptide/neuropeptide Y (AgRP/NPY) neurons. Conversely, these neurons affect Kiss1 neuron activity. NPY can suppresses firing of ARH *Kiss1* neurons kisspeptin neurons in both sexes [21], and stimulation of AgRP fibers causes a direct inhibition of Kiss1 neurons in the ARH and AVPV in mice [22]. Kiss1 neurons appear to mediate the stimulatory effects of the POMC product α-MSH on luteinizing hormone (LH) secretion [23].

These functional connections likely allow *Kiss1* neurons to coordinate reproduction and energy balance [24; 25].

*Kiss1*-expressing neurons respond to circulating metabolic cues. Some evidence suggests that leptin acts via *Kiss1*-expressing cells to exert its effects on fertility [26], although deletion of leptin receptor from Kiss*1*-expressing cells did not change puberty or fertility in mice of both sexes [27]. Insulin is another circulating factor that may influence *Kiss1*-expressing neurons [28; 29]. Only 5% of *Kiss1*-expressing neurons in all hypothalamic sites express IR protein or activated downstream signaling pathways in response to insulin [28–30]. On the other hand, IR mRNA co-expression was nearly 22% in the ARH [29]. We and others have reported a delay in the initiation of puberty in both sexes [29] or an alteration in the regulation of fasting glucose levels in male mice [30] as a result of loss of the IR in *Kiss1*-expressing cells. However, insulin sensing by *Kiss1*-expressing cells is not required for adult fertility, body growth or energy balance [29]. To explore whether insulin and leptin signaling interact, we have previously ablated both leptin and insulin signaling in *Kiss1*-expressing cells [31]. We found that the addition of leptin insensitivity did not alter the reproductive phenotype of mice with single deletion of IR; thus, *Kiss1* neurons do not directly mediate the critical role that insulin and leptin play in reproduction [31]. These studies suggest that *Kiss1*-expressing cells respond to other metabolic cues. Notably, IGF-1 and insulin act through related tyrosine kinase receptors whose signals converge on IR substrate proteins [32] and activate phosphatidylinositol 3-kinase to promote AKT signaling [33]. IGF1R and IR signaling in *Kiss1*-expressing cells may therefore jointly regulate puberty, fertility, and metabolic functions, potentially suggesting that the effects of loss of IR signaling in *Kiss1*-expressing cells may be compensated by the existence of IGF1R signaling.

IGF1R signaling plays a critical role in the regulation of several important physiological functions [34–37]. Over-expression central IGF-1 or intracerebroventricular injection of IGF-1, increases appetite, improves glucose tolerance and insulin sensitivity, and elevates UCP1 expression in BAT [34]. In addition, IGF-1 acts centrally to mediate many of its effects on reproduction [35; 36]. Notably, brain IGF1R knockout mice suffered from growth retardation, infertility, and abnormal behavior [37], while deletion of IGF1R in GnRH neurons did not impact fertility [38]. Thus, *Kiss1*-expressing neurons may mediate central IGF1R’s reproductive effects. In addition, it is currently unclear whether *Kiss1*-expressing cells in the periphery mediate the effects of IGF1 on fertility and energy use. To begin to answer these questions, we characterized the metabolic and reproductive functions of mice lacking only IGF1R (IGF1R^Kiss1^ mice) or both IGF1R and IR specifically in *Kiss1*-expressing cells (IGF1R/IR^Kiss1^ mice).

## 2. MATERIALS AND METHODS

### 2.1 Animals and genotyping

To generate mice with the IGF1R specifically deleted in *Kiss1*-expressing cells, Kiss-cre mice [39] (RRID:IMSR_JAX:023426) were crossed with IGF1R-floxed mice (RRID:IMSR_JAX:016831) and bred to homozygosity for the floxed allele. The IGF1R^loxp/loxp^ mice were designed with loxP sites flanking exon 4. Excision of exon 4 in the presence of Cre recombinase results in a truncated protein that has no functional capacities [40]. Littermates carrying only loxP sites were used as controls. To generate IGF1R/IR^Kiss1^ mice, Kiss-cre mice were crossed with IGF1R-floxed and IR floxed mice [41] (RRID:IMSR_JAX:006955) and bred to homozygosity for the floxed allele*s*. All mice were on a C57BL/6 background. Where specified, the mice also carried the Gt(ROSA)26Sor locus-inserted enhanced green fluorescent protein gene [B6.129-Gt(ROSA)26Sor^tm2Sho^/J; The Jackson Laboratory, Bar Harbor, Maine], as a reporter expressed under the control of Cre recombinase.

Mice were housed in the University of Toledo College of Medicine animal facility at 22°C to 24°C on a 12-hour light/12-hour dark cycle. They were fed standard rodent chow (2016 Teklad Global 16% Protein Rodent Diet, 12% fat by calories; Harlan Laboratories, Indianapolis, Indiana). On postnatal day (PND) 21, mice were weaned. At the end of the study, all animals were sacrificed by CO_2_ asphyxiation. Mice were genotyped using the primer pairs described in **Table 1**. We performed PCR amplification of the IGF-1 receptor floxed (flanked by *loxP* sites) genomic regions and Cre transgene in tail-derived DNA (Denville DirectAmp^TM^ Genomic DNA Amplication Kit). Additional amplification of the IR floxed genomic regions was performed by Transnetyx, Inc (Cordova, Tennessee) using a real-time PCR-based approach. The University of Toledo College of Medicine Institutional Animal Care and Use Committee approved all procedures.

**Table 1.**
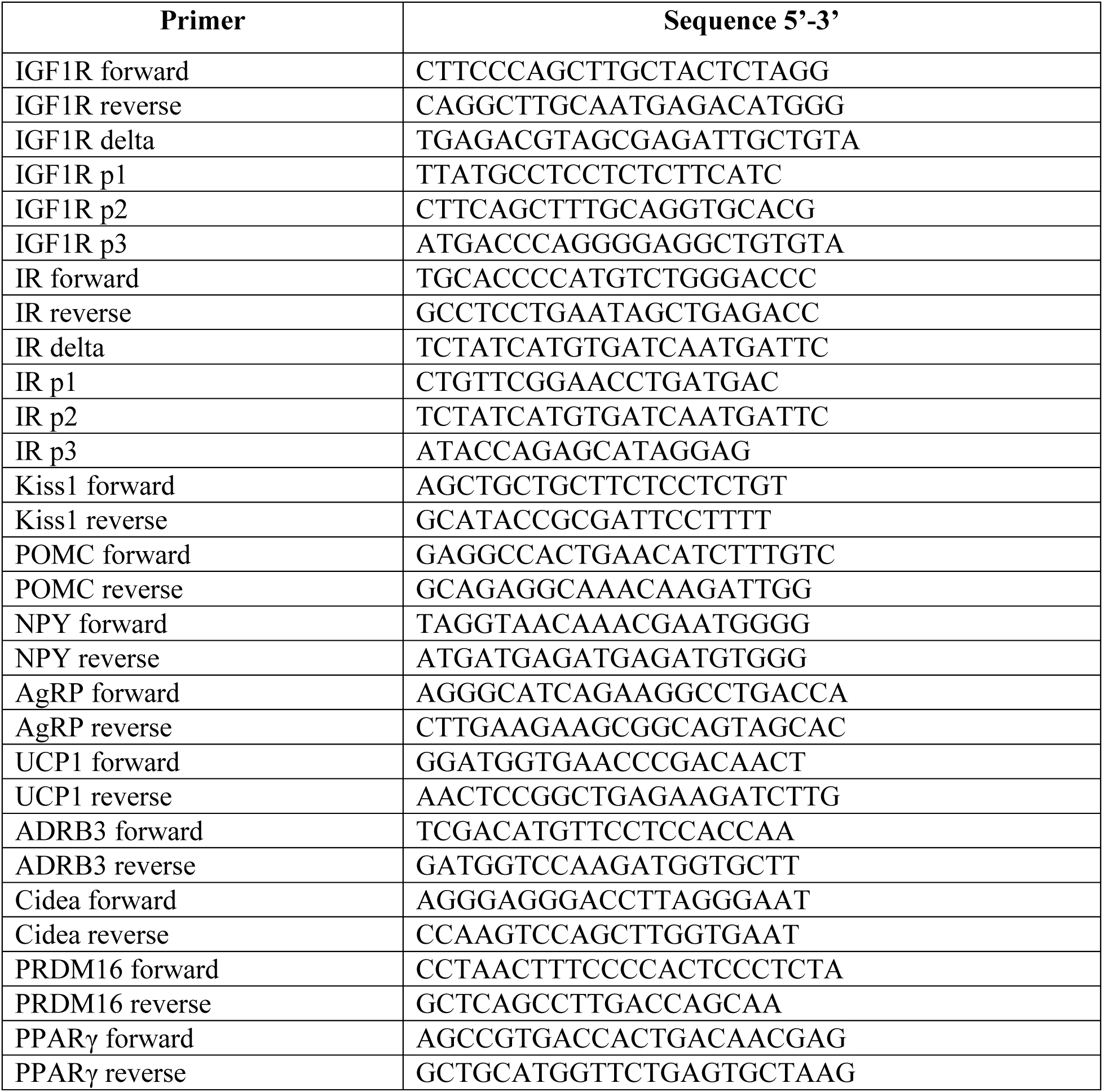
Mouse primers for measurement of gene expression.

### 2.2 Puberty and reproductive phenotype assessment

Timing of pubertal development was checked daily after weaning by determining vaginal opening (VO) in female mice or balanopreputial separation (BPS) in male mice. Cycle stages were assessed from cytology observed in saline vaginal lavages collected from females approximately 6 to 10-weeks old. The vaginal lavages evaluation was as described previously [42; 43]. First estrus age was defined as the first appearance of two consecutive days with keratinized cells after two previous days with leukocytes [42]. At 3 to 4 months of age, we examined adult fertility in female mice, who were paired with fertile adult male wild-type breeders for eight nights to determine pregnancy rate, interval from mating to birth, and litter size.

Balanopreputial separation (BPS) was checked daily from weaning by manually retracting the prepuce with gentle pressure [29]. After 3 to 4 months of age, each male mouse was paired with one fertile wild-type female for 8 nights. The paired mice were then separated, and pregnancy rate, litter size, and interval from mating to birth were recorded.

### 2.3 Tissue collection and histology

Ovaries, testes, visceral white adipose tissue (WAT) and BAT were collected and fixed immediately in 10% formalin overnight and then transferred to 70% ethanol. Tissues were embedded in paraffin and cut into 5- to 8-µm sections. Sections were stained by hematoxylin and eosin and then analyzed. For follicle and sperm quantification, a minimum of four ovaries and testes for each genotype at 5-month-old age were collected. Follicles were classified into the following categories: primordial, primary, secondary, Graafian. Testes sections were analyzed by evaluating sperm stages, including counting the number of spermatogonium, spermatocytes, spermatid, and spermatozoa using light microscopy under 20x magnification [44]. Sperm counts are reported per seminiferous tubule cross-section.

### 2.4 Quantitative real-time PCR

Hypothalamus and BAT were removed after the control, IGF1R^Kiss1^ and IGF1R/IR^Kiss1^ mice were sacrificed. Total hypothalamic and BAT RNA was extracted from dissected tissues using a RNeasy Lipid Tissue Mini Kit (QIAGEN, Valencia, California). BAT RNA was extracted by using TRIzol (Sigma-Aldrich, St. Louis, MO, USA) as described previously [45]. Single-strand cDNA was synthesized with a High-Capacity cDNA Reverse Transcription kit (Applied Biosystems) using random hexamers as primers (Appendix A). Each sample was analyzed in duplicate to measure gene expression level. A 25µM cDNA template was used in a 25µl system in 96-well plates with SYBR Green qPCR SuperMix/ROX (Smart Bioscience Inc, Maumee, Ohio). The reactions were run in an ABI PRISM 7000 sequence detection system (PE Applied Biosystems, Foster City, California). Alternatively, a 10µMcDNA template was used in a 10 µl system in 384-well plates with SYBR Green qPCR SuperMix/ROX (Smart Bioscience Inc, Maumee, Ohio). These reactions were run in a ThermoFisher QuantStudio 5 Real-Time PCR system (Applied Biosystems, Foster City, California). All data were analyzed using the comparative Ct method (2^-ΔΔCt^) with glyceraldehyde-3-phosphate dehydrogenase (GAPDH) as the housekeeping gene. GAPDH Ct values were statistically similar between groups. Relative gene copy number was calculated as 2^-ΔΔCt^ and presented as fold change of the relative mRNA expression of the control group.

### 2.5 Hormone assays

Submandibular blood was collected from 3- to 4-week-old mice (before VO or balanopreputial separation) and 3-month-old male and female mice on diestrus at 9:00 to 11:00 AM to detect LH, follicle-stimulating hormone (FSH), and estradiol levels. LH and FSH were measured via RIA performed by the University of Virginia Center for Research in Reproduction Ligand Assay and Analysis Core (Charlottesville, VA). The assay for LH had a detection sensitivity of 3.28 pg/ml, and its intra-assay and inter-assay coefficients of variance (CVs) were 4.0% and 8.6%. The detection limit of FSH assay was 7.62 pg/ml, while its intra-assay and inter-assay CVs were 7.4% and 9.1%. Serum estradiol was measured by ELISA (Calbiotech, Spring Valley, California) with a sensitivity of 3 pg/mL and intra-assay and inter-assay CVs of <10%. Testosterone in serum was measured by ELISA (Calbiotech, Spring Valley, California) with a sensitivity of 0.1 ng/mL and intra-assay and inter-assay CVs of <10%. Serum IGF-1 was measured by ELISA (Crystal Chem, Elk Grove Village, IL) with sensitivity of 0.5 to 18 ng/mL and precision intra-assay and inter-assay CVs of <10%. Serum insulin was measured by ELISA (ALPCO Diagnostics, Salem, NH) with a sensitivity of 3.0- to 200 µIU/mL and precision intra-assay and inter-assay CVs of <10%. Serum C-peptide was measured by ELISA (Crystal Chem, Elk Grove Village, IL) with sensitivity of 0.37 to 15 ng/mL and precision intra-assay and inter-assay CVs of <10%.

### 2.6 Perfusion and immunohistochemistry

3 to 6-month-old adult male mice and female mice in diestrus were deeply anesthetized by ketamine and xylazine. After brief perfusion with a saline rinse, mice were perfused transcardially with 10% formalin for 10 minutes, and the brain was removed. The brain was post-fixed in 10% formalin at 4°C overnight and immersed in 10%, 20%, and 30% sucrose at 4°C for 24 hours each. Then, 30-µm sections were cut by a sliding microtome into five equal serial sections. After rinsing in PBS, sections were blocked for 2 hours in PBS-T (PBS, Triton X-100, and 10% normal horse serum). Then, samples were incubated for 48 hours at 4°C in PBS-T-containing rabbit anti-IGF1R β antibody (1:1000; Cell signaling, Cat#9750), which has been previously validated [46]. After three washes in PBS, sections were incubated in PBS-T (Triton X-100 and 10% horse serum) containing secondary antibodies for 2 hours at room temperature. Secondary antibody was Alexa Fluor 488 (1:1,000, Thermofisher Scientific, Cat. #A-21206). Finally, sections were washed, mounted on slides, cleared, and coverslipped with fluorescence mounting medium containing DAPI (Vectasheild, Vector laboratories, Inc. Burlingame, California). Neurons were quantified in two rostrocaudal levels of the ARH (relative to bregma −1.70 and −1.94 mm). Bregma was determined according to the Allen Brain Atlas (http://mouse.brain-map.org). The quantification was performed on one side of a representative rostrocaudal level of each area analyzed.

### 2.7 Statistical analysis

Data are presented as means ± SEM. Normality testing was used to determine the normal distribution of data. If the data followed a normal distribution, One-way ANOVA was used as the main statistical method to compare the three groups, followed by the Tukey multiple comparison test. If the data did not follow a normal distribution, the Kruskal-Wallis test was used. For body weight, body length, energy expenditure, locomotor activity, glucose tolerance test (GTT), and insulin tolerance test (ITT), a two-way ANOVA was used to compare changes over time between three groups. Bonferroni multiple comparison tests were then performed to compare differences among groups. A value of *P* ≤ 0.05 was considered significant.

## 3. RESULTS

### 3.1 Disruption of IGF1R in IGF1R^Kiss1^ and IGF1R/IR ^Kiss1^ mice

To verify gene excision in *Kiss1*-expressing cells, PCR was performed on DNA from different tissues. Using this Kiss-cre line crossed with IR^loxp^ mice, our lab has previously shown excision of the floxed portion of the insulin receptor gene in the hypothalamus, as well as in the gonads, liver, and pancreas, with no ablation in the heart, gonadal fat, spleen, or muscle [29]. A 390-bp band showing IGF1R gene deletion was produced from hypothalamic tissue but appeared to be absent from ovary, liver, and fat (Figure 1A and B). To examine IGF1R expression in *Kiss1*-expressing cells in the hypothalamus, we performed double label immunohistochemistry using antibodies specific for GFP and IGF1R in *Kiss1*-cre mice crossed with a GFP reporter mouse line (Figure 1C). Approximately 20% of GFP labeled *Kiss1* cells in the ARH expressed IGF1R protein, while no double labeling was seen in the RP3V. Compared to control mice, colocalization was sharply lower in IGF1R^Kiss1^ mice (Figure 1D). No changes were seen in hypothalamic *Kiss1* mRNA expression among all groups (Figure 1E).

**Figure 1.**
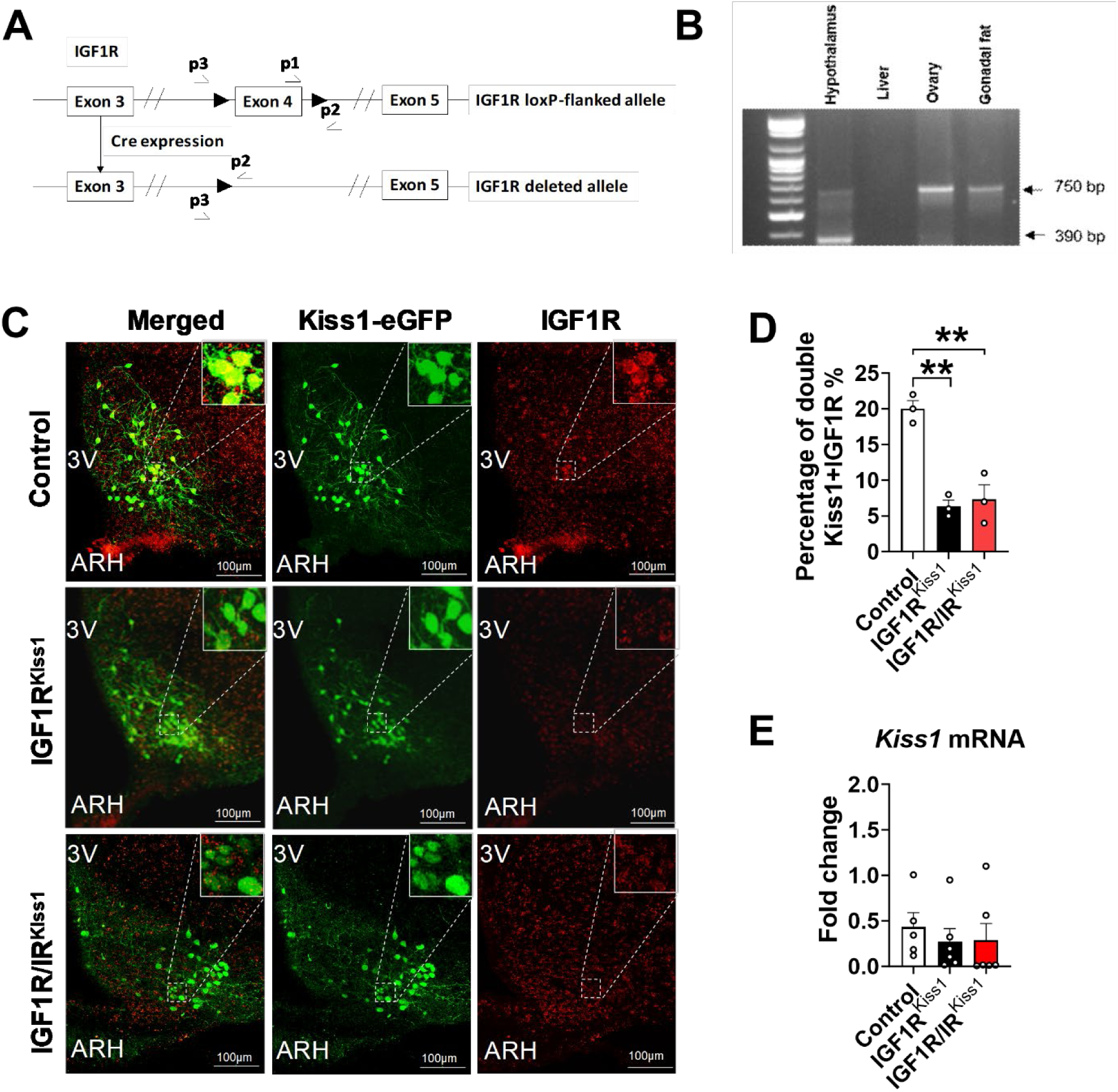
Loss of hypothalamic IGF1R and IR protein expression in Kiss1 neurons of IGF1R^Kiss1^ mice. (A) The IGF1R gene with exon 4 flanked by loxP sites (large arrows) before (top panel) and after (bottom panel) Cre recombination. Primers used for detection of the truncated IGF1R gene are labeled p1, p2, and p3. (B) PCR analysis of DNA from hypothalamus, liver, ovary, and testis of a IGF1R^Kiss1^ mouse. The excised IGF1R gene appears as a 390-bp band and the unexercised IGF1R gene sequence as a 750-bp band. (C) Colocation of *Kiss1* neuron and IGF1Rs from control and IGF1R^Kiss1^ mice. (D) Quantification of colocalization. (E) *Kiss1* mRNA expression in hypothalamus. Values throughout figure are means ±SEM. For entire figure, *p < 0.05, **p < 0.01, determined by Tukey’s post hoc test following one-way ANOVA.

### 3.2 Body growth in IGF1R^Kiss1^ and IGF1R/IR^Kiss1^ mice

Female IGF1R^Kiss1^ mice showed stunted body length growth starting from the age of weaning (Figure 2A) compared to controls. Female IGF1R/IR^Kiss1^ mice had a growth trajectory similar to the single knockout line, but neither exhibited a change in serum IGF-1 level compared to controls (Figure 2B). Male IGF1R^Kiss1^ mice exhibited reduced body length at older ages compared to controls (starting at 4 months of age), with a similar trend in the double knockout (Figure 2C). Again, no difference was seen in circulating IGF-1 levels (Figure 2D).

**Figure 2.**
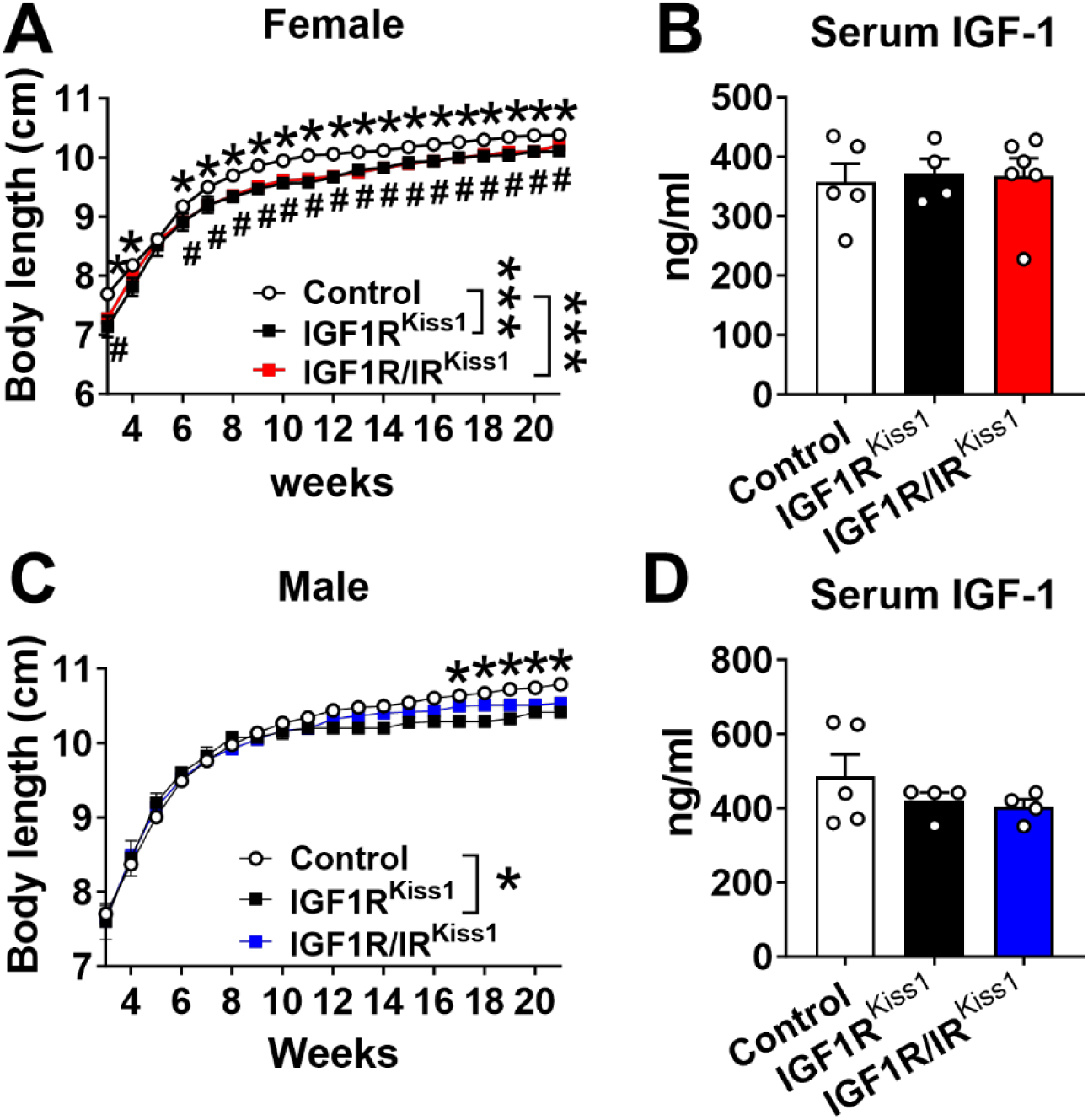
Body length and IGF-1 levels in IGF1R^Kiss1^ and IGF1R/IR^Kiss1^ Mice. (A) Body length curves and (B) serum IGF-1 levels in female control, IGF1R^Kiss1^ and IGF1R/IR^Kiss1^ mice. (C) Body length curves and (D) serum IGF-1 levels in male control, IGF1R^Kiss1^ and IGF1R/IR^Kiss1^ mice. All data are shown as means ±SEM with individual values. For the entire figure, *p < 0.05 (control vs IGF1R^Kiss1^ or IGF1R/IR^Kiss1^ mice), ^#^p < 0.05 (control vs. IGF1R/IR^Kiss1^ mice), as determined by Bonferroni’s Multiple Comparison Test following two-way ANOVA for each time point in A and C, or Tukey’s post hoc test following one-way ANOVA for B and D.

### 3.3 Altered energy balance in IGF1R^Kiss1^ and IGF1R/IR^Kiss1^ mice

Female IGF1R^Kiss1^ mice were lighter compared to controls (Figure 3A). Percentage body composition of fat and lean tissue (Figure 3B) was similar to controls, as were the absolute values of fat and lean mass (Figure 3C).

**Figure 3.**
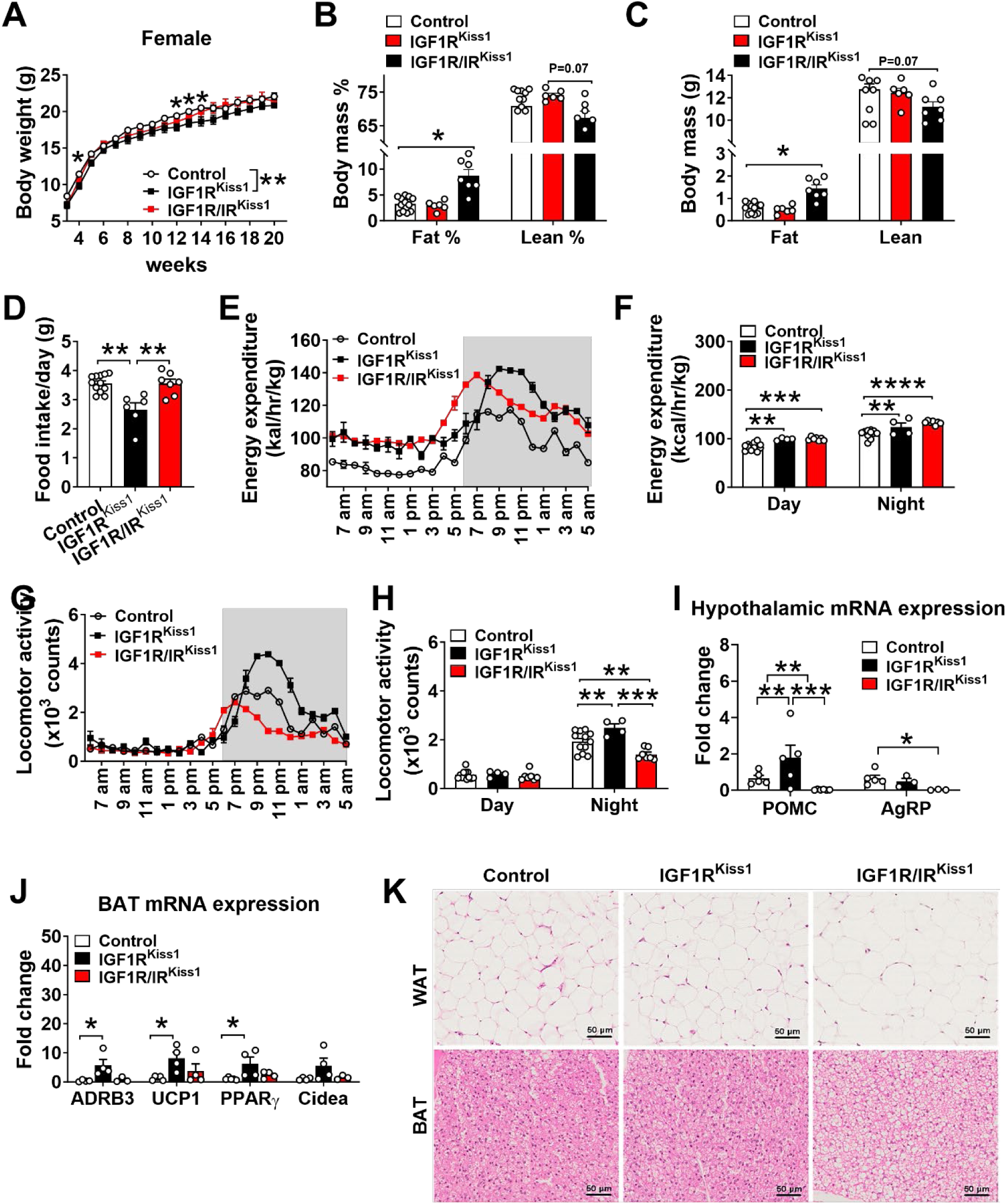
Altered energy balance in female IGF1R^Kiss1^ and IGF1R/IR^Kiss1^ mice. (A) Body weight curves, (B-C) fat and lean mass percentage and weight, and (D) food intake of female control, IGF1R^Kiss1^ and IGF1R/IR^Kiss1^ mice. (E-F) Energy expenditure and (G-H) physical activity in 4-month-old female control, IGF1R^Kiss1^ and IGF1R/IR^Kiss1^ mice. (I) Relative expression of POMC and AgRP mRNA in hypothalamus as measured by quantitative PCR in female control, IGF1R^Kiss1^ and IGF1R/IR^Kiss1^ mice. (J) Relative expression of thermogenesis markers in BAT as measured by quantitative PCR in female control, IGF1R^Kiss1^ and IGF1R/IR^Kiss1^ mice. (K) Representative sliced and HE-stained paraffin-embedded white and brown adipose tissue in 5-month-old male control, IGF1R^Kiss1^ and IGF1R/IR^Kiss1^ mice. N=4-12 per genotype. All data are shown as means ±SEM with individual values. For entire figure, *p < 0.05, **p < 0.01, ***p < 0.0001, ****p< 0.00001, determined by Bonferroni’s Multiple Comparison Test following two-way ANOVA for each time point in A, E, G; or Tukey’s post hoc test following one-way ANOVA.

In general, calorie needs correlate positively with height/length. Female IGF1R^Kiss1^ mice had lower food intake (Figure 3D) and higher energy expenditure (Figure 3E-F) and physical activity (Figure 3G-H). These findings suggest that loss of IGF1R in *Kiss1*-expressing cells led to a leaner phenotype and a tendency toward lower body weights. For mechanistic insight, we measured anorexigenic and orexigenic neuropeptide POMC, AgRP mRNA expression in the hypothalamus. POMC mRNA was higher in female IGF1R^Kiss1^ mice (Figure 3I), in alignment with decreased food intake and improved energy balance. No changes were seen in the AgRP mRNA expression (Figure 3I). We also assessed the expression of genes related to BAT thermogenesis. Female IGF1R^Kiss1^ mice had more mRNA expression of adrenoceptor beta 3 (ADRB3), uncoupling protein 1 (UCP1), and peroxisome proliferator-activated receptor gamma (PPARγ) (Figure 3J). Cidea (Figure 3J) expression did not differ. These results indicate that *Kiss1*-expressing cells regulate energy balance in response to IGF1 in a manner involving POMC neurons and activation of BAT thermogenesis.

While female IGF1R/IR^Kiss1^ mice did not show changes in body weight or lean mass (Figure 3A-C), simultaneous loss of both IGF1R and IR in *Kiss1*-expressing cells caused a nearly three-fold increase in fat mass (Figure 3B). In contrast, loss of IR [29] or IGF1R alone from *Kiss1*-expressing cells did not change body composition in female mice. These findings suggest that IGF1R and IR jointly regulate body fat mass via *Kiss1*-expressing cells in female mice. The additional deletion of IR in *Kiss1*-expressing cells reversed the lean phenotype seen in female IGF1R^Kiss1^ related to food intake, physical activity, and brown fat activation (Figure 3D, G-H, K), although energy expenditure remained elevated. In addition, hypothalamic mRNA expression of POMC and AgRP was lower in female IGF1R/IR^Kiss1^ mice than other groups (Figure 3I). These results show that IGF1R and IR signaling in *Kiss1*-expressing cells have divergent roles in regulating food intake and physical activity in female mice.

Male IGF1R^Kiss1^ mice had lower body weights prior to 6 weeks of age and inconsistently lower body weights in adulthood (Figure 4A) with no change in fat mass and lean mass percentage or absolute value (Figure 4B-C). Male IGF1R^Kiss1^ mice showed a trend toward lower food intake (Figure 4D). Unlike females, male IGF1R^Kiss1^ mice had comparable energy expenditure (Figure 4E-F) and lower physical activity (Figure 4G-H). No change was seen in BAT weight (Figure 4I). Male IGF1R/IR^Kiss1^ mice did not exhibit different body weight compared to control mice (Figure 4A), although they showed a higher fat to lean mass ratio (Figure 4B-C). The lack of IR [29] or IGF1R in *Kiss1*-expressing cells did not change body composition in males. Consistent with females, these findings suggest that *Kiss1* IGF1R and IR also play opposing roles in regulating body fat and lean mass composition in male mice. Male IGF1R/IR^Kiss1^ mice had lower energy expenditure and locomotor activity in the dark phase compared to IGF1R^Kiss1^ mice and controls, showing that both IGF1R and IR signaling in *Kiss1*-expressing cells contribute to regulating energy expenditure (Figure 4E-H). Thus, both IGF1R and IR in *Kiss1*-expressing cells are important in maintaining normal energy balance, and their effects are sex-dependent.

**Figure 4.**
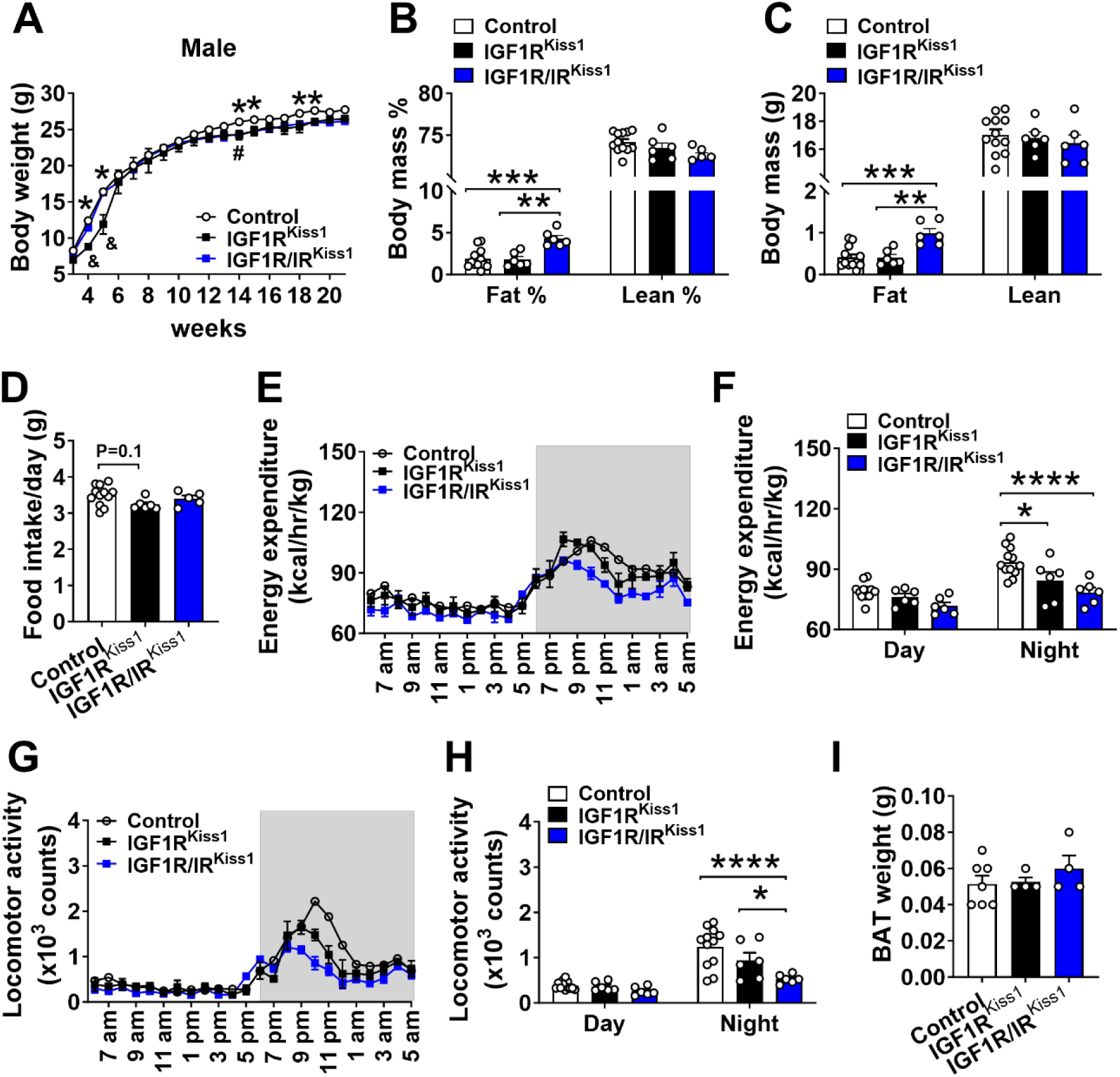
Altered energy balance in male IGF1R^Kiss1^ and IGF1R/IR^Kiss1^ mice. (A) Body weight curves, (B-C) fat and lean body mass percentage and weight, and (D) food intake of male control, IGF1R^Kiss1^ and IGF1R/IR^Kiss1^ mice. (E-F) Energy expenditure and (G-H) physical activity in 4-month-old male control, IGF1R^Kiss1^ and IGF1R/IR^Kiss1^ mice. (I) BAT weight in 5-month-old male control, IGF1R^Kiss1^ and IGF1R/IR^Kiss1^ mice. N=4-12 per genotype. All data are shown as means ±SEM with individual values. For entire figure, *p < 0.05, **p < 0.01, ***p < 0.0001, ****p< 0.00001, determined by Bonferroni’s Multiple Comparison Test following two-way ANOVA for each time point in A, E, G; or Tukey’s post hoc test following one-way ANOVA.

### 3.4 Disrupted glucose homeostasis in female and male IGF1R/IR^Kiss1^ mice

Glucose homeostasis was evaluated in 3-month-old mice. Female IGF1R^Kiss1^ mice had normal glucose regulation as shown by a GTT and ITT (Figure 5A-D). Similarly, we showed the lack of IR in *Kiss1*-expressing cells did not influence glucose homeostasis [29]. Notably, Female IGF1R/IR^Kiss1^ mice had insulin insensitivity compared to controls (Figure 5C-D), suggesting each receptor can compensate for the other’s loss. No changes were seen in fasting glucose, serum levels of insulin, C-peptide, or insulin/C-peptide ratio (Figure 5E-H). As in females, the loss of IGF1R alone in male *Kiss1*-expressing cells did not change glucose homeostasis (Figure 5I-P). Likewise, the lack of IR alone in *Kiss1*-expressing cells did not influence glucose homeostasis in male mice [29]. Importantly, male IGF1R/IR^Kiss1^ mice exhibited glucose intolerance compared to controls (Figure 5I-J). Thus, IGF1R alone in *Kiss1*-expressing cells does not control glucose homeostasis, but IGF1R and IR may cooperatively regulate glucose homeostasis in mice of both sexes.

**Figure 5.**
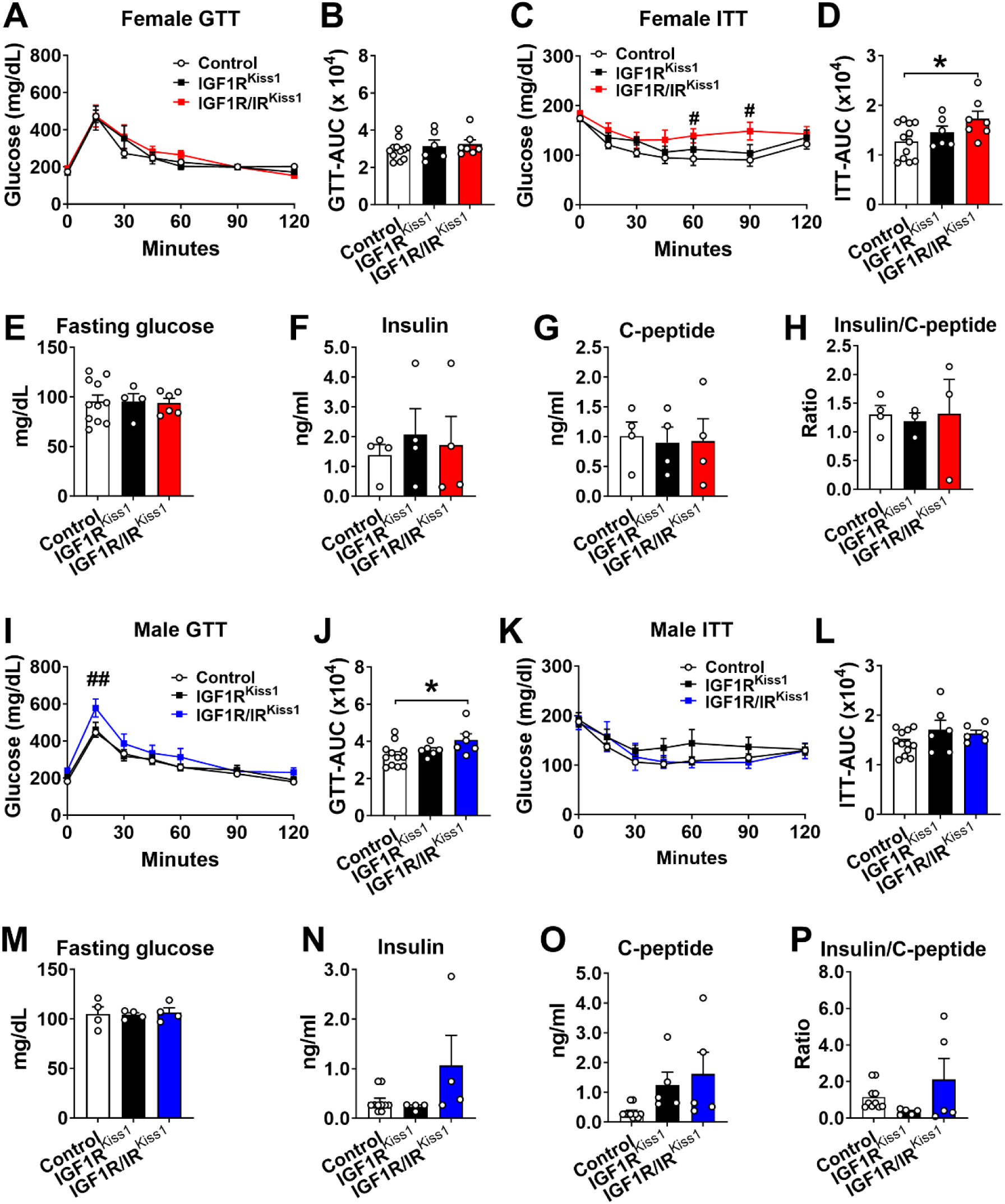
Insulin insensitivity in female IGF1R/IR^Kiss1^ mice and glucose intolerance in male IGF1R/IR^Kiss1^ mice. (A) Glucose tolerance test (GTT), (B) area under curve (AUC) of GTT (GTT-AUC), (C) insulin tolerance test (ITT) (D) and AUC of ITT (ITT-AUC) in 3-month-old female control, IGF1R^Kiss1^ and IGF1R/IR^Kiss1^ mice. (E) Fasting glucose, (F) insulin, (G) C-peptide, and (H) insulin: C-peptide ratio in 3-month-old female control, IGF1R^Kiss1^ and IGF1R/IR^Kiss1^ mice. (I) GTT, (J) GTT-AUC, (K) ITT, and (L) ITT-AUC in 3-month-old male control, IGF1R^Kiss1^ and IGF1R/IR^Kiss1^ mice. (M) Fasting glucose, (N) insulin, (O) C-peptide, and (P) insulin: C-peptide ratio in 3-month-old male control, IGF1R^Kiss1^ and IGF1R/IR^Kiss1^ mice. N=6-12 per genotype. All data are shown as means ±SEM with individual values. For entire figure, *p < 0.05, ^#^p < 0.05 (control vs IGF1R/IR^Kiss1^ mice), determined by Bonferroni’s Multiple Comparison Test following two-way ANOVA for each time point in A, C, I and K; or Tukey’s post hoc test following one-way ANOVA.

### 3.5 Delayed pubertal development in female and male IGF1R^Kiss1^ and IGF1R/IR^Kiss1^ mice

To evaluate the pubertal development in female mice, we examined VO and the timing of the first entrance into estrus. Female IR^Kiss1^ mice experienced a delay in both parameters [29]. Here we found that female IGF1R^Kiss1^ mice experienced VO approximately 3 days later (Figure 6A) and first estrus approximately 4 days later than controls (Figure 6B). No differences were seen in serum LH levels (Figure 6C) between female control and IGF1R^Kiss1^ mice at 4-weeks of age. However, female IGF1R^Kiss1^ mice showed significantly but transiently higher levels of FSH than controls (Figure 6D). Female IGF1R/IR^Kiss1^ mice also had delayed VO age and first estrus age, but IGF1R/IR^Kiss1^ and IGF1R^Kiss1^ mice were comparable (Figure 6A-D).

**Figure 6.**
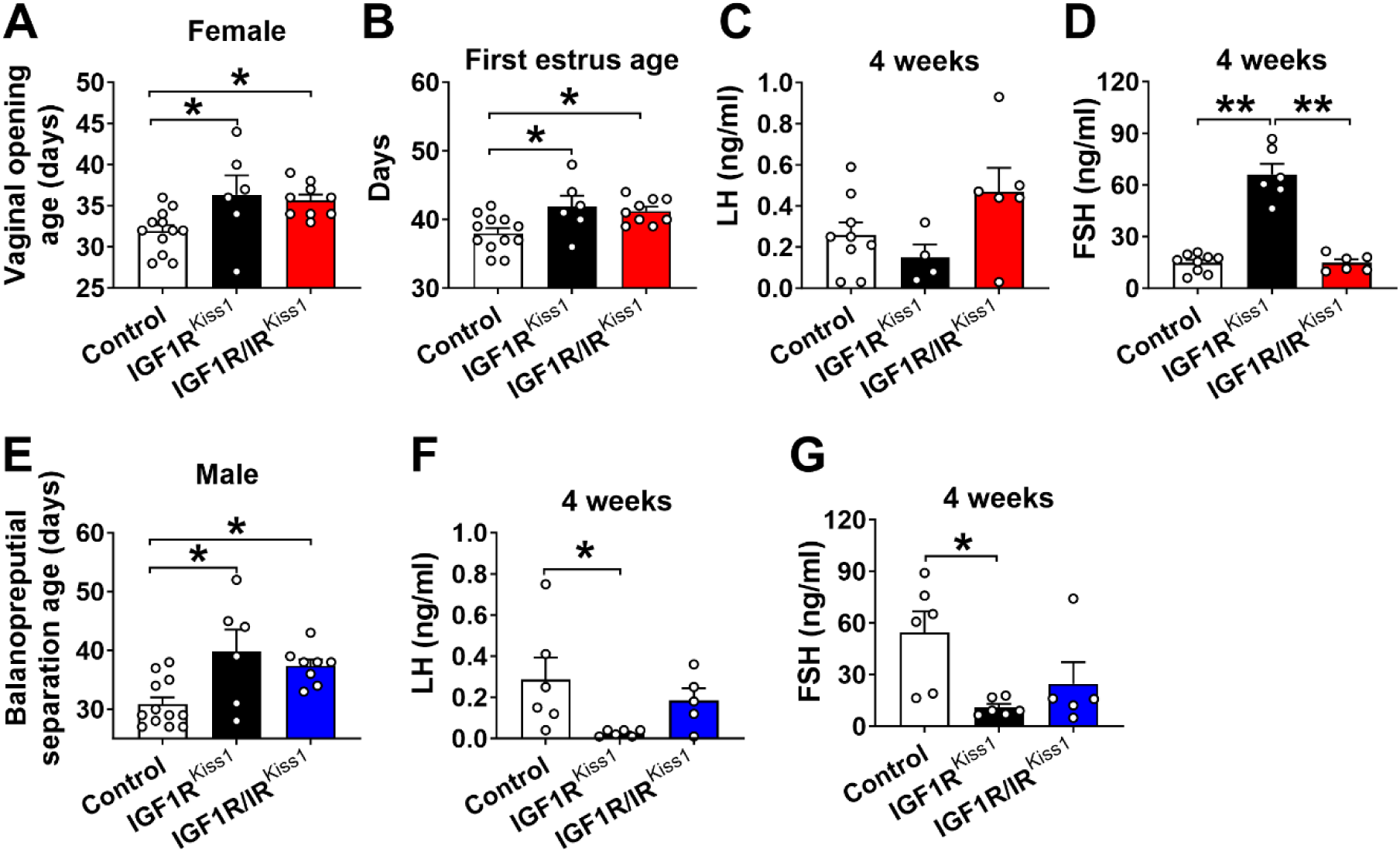
Delayed pubertal development in IGF1R^Kiss1^ and IGF1R/IR^Kiss1^ mice. (A) Vaginal opening age and (B) first estrus age in female control, IGF1R^Kiss1^ and IGF1R/IR^Kiss1^ mice. (C-D) Serum levels of LH and FSH in 4 weeks-old female control, IGF1R^Kiss1^, and IGF1R/IR^Kiss1^ mice. (E) Balanopreputial separation age and serum levels of (F) LH and (G) FSH in male control, IGF1R^Kiss1^, and IGF1R/IR^Kiss1^ mice. N=5-12 per genotype. LH luteinizing hormone, FSH follicle-stimulating hormone. All data are shown as means ±SEM with individual values. For entire figure, *\**p < 0.05, **p < 0.01, determined by Tukey’s post hoc test following one-way ANOVA.

Balanopreputial separation (BPS) was evaluated as an external sign of puberty in male rodents. Male IGF1R^Kiss1^ mice displayed delayed pubertal development (Figure 6E) associated with lower LH and FSH levels at 4-weeks of age compared to controls (Figure 6F-G). Male IGF1R/IR^Kiss1^ mice had delayed pubertal development (Figure 6G), but there was no difference between IGF1R^Kiss1^ and IGF1R/IR^Kiss1^ mice. These findings suggest that IGF1R in *Kiss1*-expressing cells is required for normal pubertal onset in both female and male mice and normal gonadotropin levels in males during the pubertal transition.

We further performed linear regression analysis to examine whether the delayed onset of puberty was associated with the lower body weight. We found that lower body weight at 3 weeks of age was associated with later onset of puberty as indicated by the VO age in female mice (Supplemental Figure 1A). However, no effects of body weight were seen on the age of pubertal completion in females, namely age of first estrus. For males, balanopreputial separation age was not associated with body weight (Supplemental Figure 1B-C). These findings suggest a particular susceptibility of pubertal initiation to low body weight in females.

### 3.6 Reproductive deficits in female and male IGF1R^Kiss1^ and IGF1R/IR^Kiss1^ mice

Adult female IGF1R^Kiss1^ and IGF1R/IR^Kiss1^ mice had normal estrus cyclicity (Figure 7). To evaluate adult reproductive function, we performed fertility testing in both female and male mice. Even though the overall pregnancy rate (Figure 8A) in all female groups was comparable, the numbers of pups per litter born to IGF1R^Kiss1^ females mated with control males (Figure 8B) was lower. No differences were seen in serum levels of LH (Figure 8C), FSH (Figure 8D), estradiol (Figure 8E), ovary and uterine weight (Figure 8F), or the numbers of follicles (Figure 7G). No differences were seen between female IGF1R/IR^Kiss1^ and IGF1R^Kiss1^ mice (Figure 8A-G), and IR^Kiss1^ mice did not show impaired fertility [29]. Our findings suggest IGF1R in *Kiss1*-expressing cells is not required for ovulation but may play a pregnancy-related role that influences litter size.

**Figure 7.**
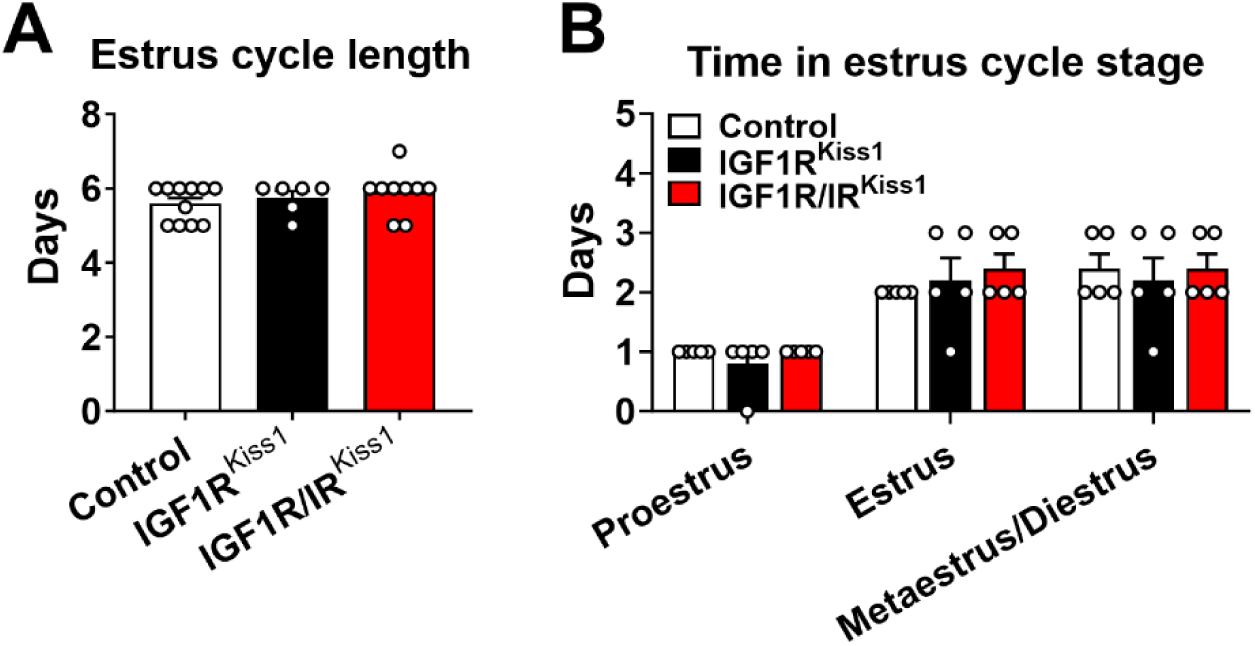
Estrus cyclicity in IGF1R^Kiss1^ and IGF1R/IR^Kiss1^ mice. (A) estrus cycle length and (B) days spent in each estrus cycle stage in female control, IGF1R^Kiss1^ and IGF1R/IR^Kiss1^ mice. N=6-12 per genotype. P proestrus, E estrus, M metestrus, D diestrus. All data are shown as means ±SEM with individual values.

**Figure 8.**
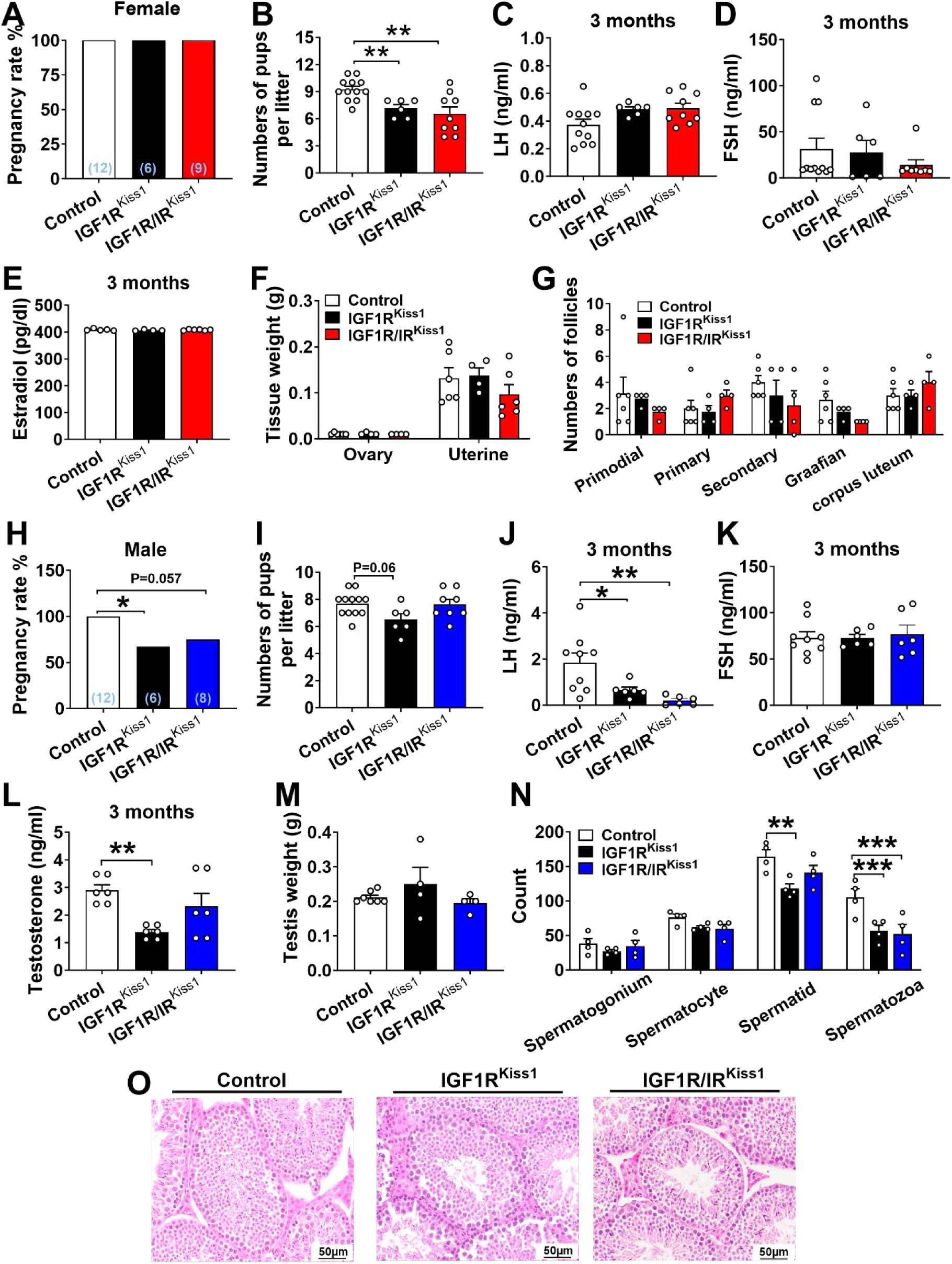
Adult fertility in IGF1R^Kiss1^ and IGF1R/IR^Kiss1^ mice. (A) Pregnancy rate, (B) numbers of pups/litter, (C) serum LH, (D) FSH and (E) estradiol levels on diestrus in 3- to 4-month-old female control, IGF1R^Kiss1^ and IGF1R/IR^Kiss1^ mice. (F) Uterine and ovary weight, and (G) ovarian follicle numbers in 5-month-old female control, IGF1R^Kiss1^ and IGF1R/IR^Kiss1^ mice. (H) Pregnancy rate, (I) numbers of pups/litter, (J) serum LH (K), LH/FSH ratio and (L) testosterone levels in 3- to 4-month-old male control, IGF1R^Kiss1^ and IGF1R/IR^Kiss1^ mice. (M) Testis weight and (N) analysis of cross-sectional testes seminiferous tubule in 5-month-old male, IGF1R^Kiss1^ and IGF1R/IR^Kiss1^ mice. (O) Representative sliced and HE-stained paraffin-embedded testes in 5-month-old male control, IGF1R^Kiss1^ and IGF1R/IR^Kiss1^ mice. N=4-12 per genotype. All data are shown as means ±SEM with individual values. For the entire figure, *p < 0.05 and **p < 0.01, determined by Tukey’s post hoc test following one-way ANOVA, except G and N determined by Bonferroni’s Multiple Comparison Test following two-way ANOVA.

Male IGF1R^Kiss1^ mice produced fewer pregnancies (Figure 8H) but similar numbers of pups per litter compared to controls (Figure 8I). Male IGF1R^Kiss1^ mice also had lower serum LH, FSH, and testosterone levels (Figure 8J-L). No change in testis weight was seen (Figure 8M). In cross-sections of the seminiferous tubule, male IGF1R^Kiss1^ mice had fewer of spermatids and spermatozoa (Figure 8N), indicating reduced spermatogenesis [47]. Male IGF1R/IR^Kiss1^ mice had similar, or slightly improved, fertility parameters compared to IGF1R^Kiss1^ mice; they tended toward a lower pregnancy rate with a lower LH and LH/FSH ratio and spermatozoa count compared to controls (Figure 8H-N). Representative histological images of seminiferous tubule are shown in Figure 8O. Consistent with females, these findings imply that IGF1R in *Kiss1*-expressing cells plays a vital role in regulating adult fertility, gonadotropins, sex hormones, and spermatogenesis in male mice.

## 4. DISCUSSION

In this study, we have explored the roles of the growth factors IGF-1 and insulin in the control of metabolism and reproduction, finding that IGF1R, but not IR, signaling in *Kiss1*-expressing cells is crucial for body growth, energy balance, normal timing of pubertal onset and male reproductive functions. Interestingly, IGF1R effects on energy balance are sex-dependent, in that female IGF1R^Kiss1^ mice had lower body weight and food intake, plus higher energy expenditure and physical activity. We also found *Kiss1* IGF1R and IR serve cooperative roles in the regulation of metabolic functions in mice.

The joint roles of IGF1R and IR in peripheral tissues, including fat and muscle, are well established [48; 49]. For example, combined deletion of IGF1R and IR from fat tissue results in cold intolerance due to impaired BAT development and function, while single receptor deletions have normal thermoregulation [48]. In muscle, IRs and IGF1Rs work together to maintain muscle growth; only when both are deleted do the mice have a substantial decrease in fiber size and muscle mass [49]. Similarly, simultaneous loss of IGF1R and IR in *Kiss1*-expressing cells caused glucose intolerance and insulin insensitivity, which was not seen in IGF1R^Kiss1^ or IR^Kiss1^ mice [29]. We previously showed that deletion of the insulin receptor (IR) alone from Kiss1 expressing cells had no effect on body weight, body fat composition, or food intake [29]. Compared to either IGF1R^Kiss1^ mice or controls, IGF1R/IR^Kiss1^ mice of both sexes exhibited markedly higher fat mass percentages. Overall, our findings identify a cooperative role for IGF1R and IR in *Kiss1*-expressing cells in the regulation of WAT formation and glucose homeostasis.

The constraints of using conditional gene deletion affect the interpretation of this study. Compensation for loss of signaling may occur during early life, possibly leading to a less pronounced phenotype not reflecting gene function in adult mice. In addition, kisspeptin expression is not limited to the brain; rather, several relevant tissues are reported to contain some kisspeptin producing cells in mice. We and others have previously found low or absent kiss1 expression in the liver, and a lack of expression in muscle, subcutaneous fat, and the gonadal fat pad in healthy mice [29; 50], but kisspeptin is reportedly produced by hepatocytes in response to glucagon stimulation and is increased in mouse models of T2DM [51]. While our lab has previously shown excision of the floxed portion of the IR gene in the liver driven by this Kiss-cre line [29], our results do not reflect the dramatic reduction in systemic insulin sensitivity that has been shown as a consequence of loss of insulin signaling in hepatocytes [52]. Further, healthy mature hepatocytes do not express appreciable levels of the IGF1 receptor [53; 54], and we were not able to detect IGF1R gene deletion in this tissue. A potential role for Kiss1 in the mouse kidney and gut is underexplored [29; 55]. The mouse pancreas, and specifically alpha and beta cells have been reported to show kisspeptin immunoreactivity [29; 56]; insulin and IGF1 signaling have important influences on gene expression, cell survival, and proliferation in these cells [57–59]. Again, the phenotype of the mice in this study is milder than would be expected with deletion of the IGF1R or both the IGF1R and IR in cells of the pancreas [59–62]. Nevertheless, we cannot exclude the possibility that Kiss1 expressing cells in these tissues contribute to the metabolic status of IGF1R/IR^Kiss1^ mice.

The mechanism leading to altered glucose and body composition regulation in IGF1R/IR^Kiss1^ mice may involve direct action on Kiss1 ARH neurons, since they perceive hormonal cues and communicate bidirectionally with neighboring neurons such as POMC neurons that influence reproduction and energy balance [26; 63]. Anorexigenic POMC neurons in the ARH are crucial for maintaining normal energy balance in both humans and rodents [64], while also influencing pubertal timing [23]. We found greater POMC mRNA expression in female IGF1R^Kiss1^ mice, which may relate to their lower food intake and body weight. Female mice have more POMC neurons and higher POMC neuronal activity than males [65], perhaps explaining why female IGF1R^Kiss1^ mice showed a more pronounced metabolic phenotype than males. Additionally, POMC neurons can regulate BAT thermogenesis through the sympathetic nervous system (SNS) [66]. Indeed, expression of BAT thermogenesis and SNS activation genes was higher in female IGF1R^Kiss1^ mice. Therefore, one plausible mechanism for the ability of IGF1R in *Kiss1*-expressing cells to modulate energy balance is through communication with POMC neurons and activation of SNS activity and BAT thermogenesis.

IGF1R^Kiss1^ mice displayed mild growth retardation in the form of a shorter body length throughout life. We were unable to detect an endocrine cause for this phenotype since there were no changes in circulating IGF-1 levels in IGF1R^Kiss1^ and IGF1R/IR^Kiss1^ mice, and limited serum sample volume prevented growth hormone (GH) measurement. Whether neuronal or non-neuronal cells are responsible for the growth deficit is unclear. Homozygous brain IGF1R knockout mice, whose IGF1Rs were depleted in neurons and glia, displayed microcephaly with severe growth retardation [37], while the heterozygous brain IGF1R knockout mice weighed 90% of wild-type controls at 3 months of age and were 5% shorter in length [37]. However, ablation of the IGF1R in somatostatin (SST) neurons, growth hormone releasing hormone (GHRH) neurons, and/or pituitary somatotrophs did not influence body growth through 14 weeks of age [67; 68]. Thus, there is a potential role for *Kiss1* neurons as targets underlying the ability of brain IGF1Rs to modulate growth. It should be noted that GH receptor ablation in kisspeptin neurons did not affect pubertal timing, estrous cycle length, body weight, or body length [69]. Interestingly, during puberty in female mice a subpopulation of ERα positive GHRH neurons become Kiss1-positive neurons, potentially permitting crosstalk between the growth and reproductive axes [70].

We found IGF1R^Kiss1^ mice displayed delayed puberty and a smaller number of pups per litter in females, while males showed delayed puberty, diminished adult fertility, and disrupted gonadotropin and testosterone levels. The low LH levels at 4 weeks and 3 months in male IGF1R^Kiss1^ mice suggest that IGF1R signaling in *Kiss1*-expressing neurons is necessary for the activation of the male HPG axis during puberty and adult fertility. Measures of adult fertility were similar between IGF1R^Kiss1^ and IGF1R/IR^Kiss1^ mice, suggesting that IGF1R plays a more dominant role in the regulation of reproduction. Some of the reproductive deficits seen in this mouse model may involve the loss of IGF1 receptors in *Kiss1*-expressing cells within the gonads or other peripheral tissues expressing insulin and IGF1 receptors that impact their function [71; 72]. We and others have reported kiss1 expression in the testis [29; 50]. Indeed, kisspeptin is expressed in mouse Leydig cells and is regulated by reproductive hormone levels [50; 73]. Thus, local impairment of Leydig cell function may contribute to reduced testosterone levels and male subfertility. Kisspeptin is also produced in ovarian granulosa cells as oocytes mature [74; 75], and may be found in the pituitary [76; 77] and uterus [78]. The current Kiss-cre line can drive excision of the floxed portion of the IR gene in the gonads [29]. The inability of IGF1 to promote expression of steroidogenic genes [79] could plausibly explain why female IGF1R^Kiss1^ mice displayed delayed puberty despite normal LH levels and high levels of FSH during the pubertal transition. However, the apparent lack of IGF1R deletion in the ovary argues against the idea that reduced ovarian hormone negative feedback elevates FSH levels in peripubertal IGF1R^Kiss1^ mice. Overall, our results confirm that factors controlling growth and metabolism, such as IGF1, regulate fertility via *Kiss1-*expressing cells. Additional research is needed to understand the contribution of central and peripheral *Kiss1*-expressing cells to metabolic factors promoting pubertal development and adult fertility.

## 5. CONCLUSION

This study assessed the roles of IGF1R and IR in *Kiss1*-expressing cells in the regulation of metabolic and reproductive functions in mice. We found IGF1R in *Kiss1*-expressing cells is necessary for the regulation of body growth and the activation of the HPG axis during puberty and adult fertility, particularly in males. IGF1R signaling in *Kiss1*-expressing cells is critical for maintaining normal metabolic functions, including food intake and energy balance, potentially through central effects on POMC signaling and BAT thermogenesis induction. Notably, our study demonstrates that IGF1R and IR signaling in *Kiss1*-expressing cells have divergent and compensatory effects in regulating fat mass composition and glucose homeostasis. These findings emphasize the tight link between the systems controlling energy balance and fertility and the critical function of metabolic factors such as IGF-1 in facilitating homeostatic communication.

## AUTHOR CONTRIBUTIONS

Conceptualization: JWH; Data Collection: MW, SMP, JD, DM; Formal Analysis: MW, JWH; Manuscript writing and editing: MW, YX, JWH; Supervision; JWH.

## ACKNOWLEDGEMENT

This work was supported by National Institutes of Health grant HD104418 to JWH and U.S. Department of Agriculture Grant 51000-064-01S to YX.

---

**Supplemental Figure 1.**
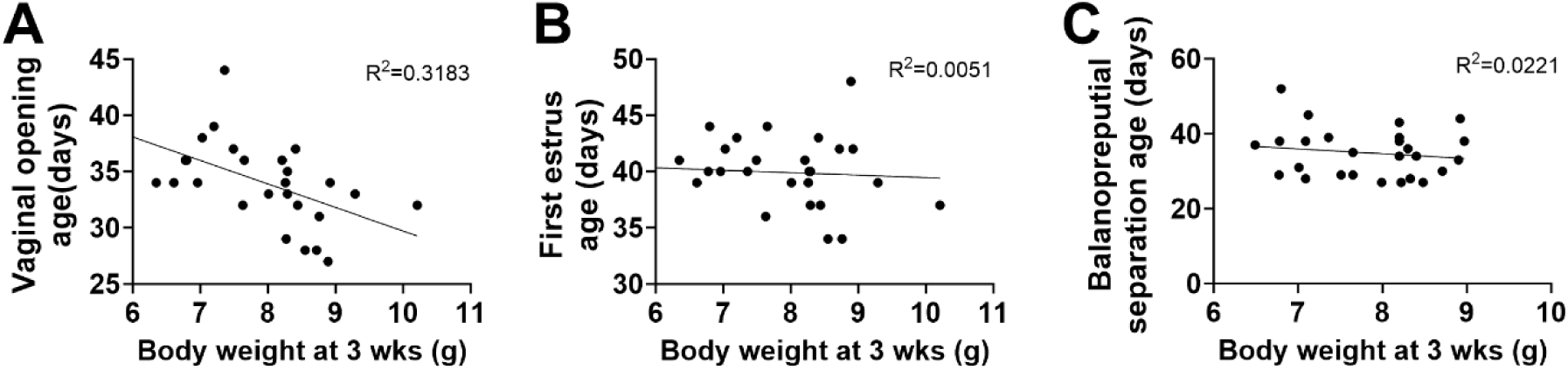
Relationship between the onset of puberty and body weight. Relationship between (A) vaginal opening age, (B) first estrus age, (C) balanopreputial separation age and body weight at 3 weeks of age. Correlation coefficient is given as a measure of linear regression between the two variables.

